# Genome-wide analysis of *cis*-regulatory changes in the metabolic adaptation of cavefish

**DOI:** 10.1101/2020.08.27.270371

**Authors:** Jaya Krishnan, Chris W. Seidel, Ning Zhang, Jake VanCampen, Robert Peuß, Shaolei Xiong, Alexander Kenzior, Hua Li, Joan W. Conaway, Nicolas Rohner

## Abstract

Changes in cis-regulatory elements play important roles in adaptation and phenotypic evolution. However, their contribution to metabolic adaptation of organisms is less understood. Here we have utilized a unique vertebrate model, *Astyanax mexicanus,* different morphotypes of which survive in nutrient-rich surface and nutrient-deprived cave water to uncover gene regulatory networks in metabolic adaptation. We performed genome-wide epigenetic profiling in the liver tissue of one surface and two independently derived cave populations. We find that many cis-regulatory elements differ in their epigenetic status/chromatin accessibility between surface fish and cavefish, while the two independently derived cave populations have evolved remarkably similar regulatory signatures. These differentially accessible regions are associated with genes of key pathways related to lipid metabolism, circadian rhythm and immune system that are known to be altered in cavefish. Using *in vitro* and *in vivo* functional testing of the candidate cis-regulatory elements, we find that genetic changes within them cause quantitative expression differences. We characterized one cis-regulatory element in the *hpdb* gene and found a genomic deletion in cavefish that abolishes binding of the transcriptional repressor IRF2 *in vitro* and derepresses enhancer activity in reporter assays. Genetic experiments further validated a cis-mediated role of the enhancer and suggest a role of this deletion in the upregulation of *hpdb* in wild cavefish populations. Selection of this mutation in multiple independent cave populations supports its importance in the adaptation to the cave environment, providing novel molecular insights into the evolutionary trade-off between loss of pigmentation and adaptation to a food-deprived cave environment.

## Introduction

Cis-regulatory elements are a major target of evolution for shaping phenotypic diversity (Long et al., 2016; Prescott et al., 2015; Wittkopp and Kalay, 2011) and helping organisms adapt to various environmental niches (Partha et al., 2017; Thompson et al., 2018). The role of cis-regulatory changes in the evolution of metabolic adaptations is less well understood. The Mexican tetra, *Astyanax mexicanus,* with its two morphotypes – the river-dwelling surface fish and the cave-dwelling cavefish, provides an exceptional model system to study evolutionary changes at a genetic and genomic level. The surface fish live in nutrient-rich rivers while the cave populations are well adapted to survive in the dark and nutrient-deprived caves, which they colonized around 150,000 years ago (Bradic et al., 2012; Herman et al., 2018). Importantly, many of the cave populations have independently adapted to different cave environments (Coghill et al., 2014). Such repeated evolution allows the study of whether the same genes and regulatory networks were utilized in adaptation or if evolution took different paths to arrive at similar phenotypes.

Missense mutations have been identified in key metabolic genes like *insra* and *mc4r* that help cavefish to survive in low-nutrient conditions (Aspiras et al., 2015; Riddle et al., 2018). However, the contribution of regulatory changes in the complex metabolic phenotypes of cavefish has not been investigated. Here, we generated a high-resolution, genome-wide map of candidate *cis*-regulatory elements in *Astyanax mexicanus*. We focused our study on the liver because of its central role in glucose metabolism including glycolysis, TCA cycle, gluconeogenesis, glycogen and lipid metabolism (Rui, 2014). Analysis of the regulatory landscapes of the two independent cave populations, Pachón and Tinaja, revealed remarkable similarity indicating signs of repeated evolution at an epigenetic level. Importantly, we identified a large number of putative *cis*-regulatory elements (CREs) that have lost or gained regulatory potential during the adaptation to the cave environment. We further show that several CREs display different epigenetic states between surface fish and cavefish which is correlated with their ability to modulate varying levels of reporter gene expression. Analyses of one of these CREs inside the *hpdb* gene, a tyrosine metabolism pathway gene, showed that deletion of a binding site for IRF2 repressor protein in cavefish is sufficient to drive increased gene expression. This suggests a potential adaptive role for the mutation, with a trade-off between the conversion of tyrosine to melanin and TCA cycle intermediates. Our study not only demonstrates the value of using independently evolved populations to identify non-coding genomic loci relevant for cave adaptation, but also provides a large number of candidates for future studies on metabolic adaptation.

## Results

### Genome-wide annotation of cis-regulatory elements in liver tissue

We analyzed accessible chromatin, histone modifications, and gene expression to generate a genome-wide epigenetic landscape in the liver tissue of surface fish and two independently derived cave morphotypes – Pachón and Tinaja (Fig. 1a; Fig. S1a) (Bradic et al., 2012; Dowling et al., 2002). We first performed ATAC-seq to map accessible chromatin regions of the genome as a basis for genome-wide identification of candidate CREs (Daugherty et al., 2017; Gross and Garrard, 1988; Klemm et al., 2019). We identified a total of 94,175 accessible chromatin regions or putative cis-regulatory elements (CREs) regions across the three populations – 68,002 in surface, 69,178 in Pachón and 73,762 in Tinaja (Supplementary Information SI-1). 19.09% of these putative CREs were in promoter regions (−2,000 to +2,000 bp from the nearest transcription start site (TSS)), 25.78% were within 10kb from the TSS, 38.29% were within 50kb from the TSS, while the remaining 16.82% were greater than 50kb away from any TSS (Fig. 1b).

**Figure 1:**
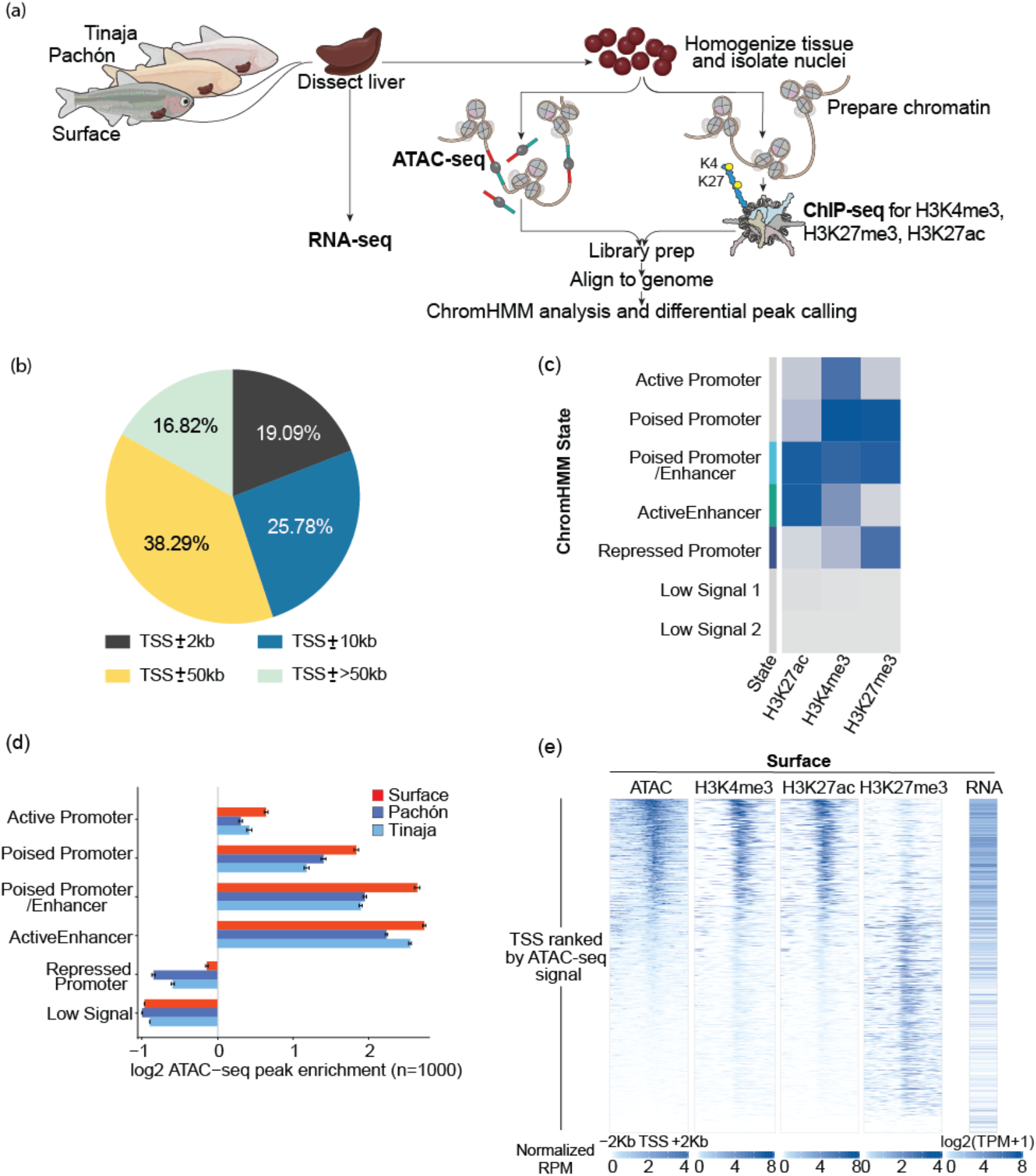
Genome-wide annotation of cis-regulatory elements. (a) Schematic of the experimental design. (b) Pie chart showing genomic distribution for the CREs mapped. We defined promoters as CREs within ±2kb of the transcription start site (TSS); Proximal CREs within ±10kb of TSS; Distal elements within ±50kb of TSS and the rest are categorized as intergenic. (c) Heatmap showing the chromatin features comprising each of the seven chromatin states into which the CREs were categorized using ChromHMM. The two low signal states belong to different genomic regions as depicted in Fig. S1b. (d) Enrichment of ATAC-seq peaks in the ChromHMM states. The two low signal states were combined into a single low signal state for better depiction. The x-axis shows the average enrichment ratio and the standard deviation (n=1000) for each ChromHMM state. (e) Heatmaps for ATAC-seq, H3K4me3, H3K27ac and H3K27me3 signal at all the promoters (TSS ±2Kb) along with the corresponding RNA levels for the surface fish. Plotted are reads per million (RPM) normalized to ChIP-seq signals at promoters (refer to Fig. S1 for cavefish heat maps).

To further characterize the accessible chromatin regions, we performed ChIP-seq for active histone marks, H3K4me3, and H3K27ac, and the repressive histone mark H3K27me3. We classified the whole genome based on histone marks using ChromHMM, an automated computational pipeline for learning chromatin states based on a Hidden Markow Model (Ernst and Kellis, 2012) (Fig. 1c; Fig. S1b). Genomic regions marked with the active histone modification H3K27ac were classified as active, with H3K27me3 mark as repressed while those marked with all three histone marks as poised (Zhou et al., 2011). In addition, active promoters were classified based on their H3K4me3 mark (reviewed in Zhou et al., 2011) (Fig. 1c; Fig. S1b). Regions with low intensity of all histone marks were classified into low signal states and combined together for further analysis (Fig. S1b). We observed that regions marked by active chromatin marks as well as poised regulatory regions had accessible chromatin while those marked solely by the repressive histone modification H3K27me3 and the regions with low signal chromatin states had relatively inaccessible chromatin (Fig. 1d). We performed bulk RNA-seq on the liver tissues and integrated gene expression data with epigenetic profile and observed that expression of genes was positively correlated with active and poised marks and inversely correlated with repressive marks (Fig. 1e; Fig. S1c, d; Supplementary Information SI-2). Together these analyses highlight that chromatin accessibility can serve as a means of identifying putative CREs that correlate with the expression of the associated genes in this system.

We further analyzed these open chromatin regions for sequence conservation across 11 fish species (Cooper et al., 2005). Regions marked by a chromatin feature were roughly 30% more conserved than randomly picked regions in the genome (Fig. S2a, b). We extended sequence conservation analyses to known regulatory regions in the human liver. We found that 94 out of 441 liver-specific human enhancers had sequence conservation and were marked as open chromatin in our dataset, indicating that some of the putative CREs we identified are conserved in vertebrates and could be involved in gene regulatory networks that control conserved metabolic processes.

### Morphotype-biased accessible chromatin regions associate with key metabolic pathways

To understand how the cis-regulatory network has evolved during cave adaptation, we focused on regions with very different chromatin accessibility between surface and cave populations. We identified 33,176 differentially accessible regions between surface and Pachón and 35,140 between surface and Tinaja (Fig. 2a). Interestingly, we identified fewer differentially accessible regions between the two cave populations, suggesting a greater divergence between the surface and cave populations than the cave populations (Fig. 2a). We further analyzed highly differentially accessible regions mapping close (<10Kb) to genes and observed that 74.4% of the regions that were accessible in surface fish (surface-biased CREs) lost accessibility in both the cave populations (Fig. 2b, c). Similarly, 77.4% of the regions that gained accessibility in Pachón as compared to surface also gained accessibility in Tinaja (cave-biased CREs) (Fig. 2d, e). These comparisons indicate that both the cave populations have gained or lost accessible chromatin states with regulatory potential in a very similar set of regions of the genome during evolution. A similar pattern was also reflected in the genome-wide pattern of histone modifications and gene expression between the two cavefish populations as compared to the surface (Fig. S2c-f). This high degree of similarity in the chromatin landscape of the two cave populations point towards modifications of existing regulatory circuitry in the surface fish and that the system is robustly wired to use similar sets of gene regulatory mechanisms in independently derived populations. Hence, to further understand the cave-adaptive cis-regulatory features, we focused on the regions that were similarly biased in both the cave populations i.e. cave-biased CREs (c-CREs) and compared with the surface-biased CREs (s-CREs) regions.

**Figure 2:**
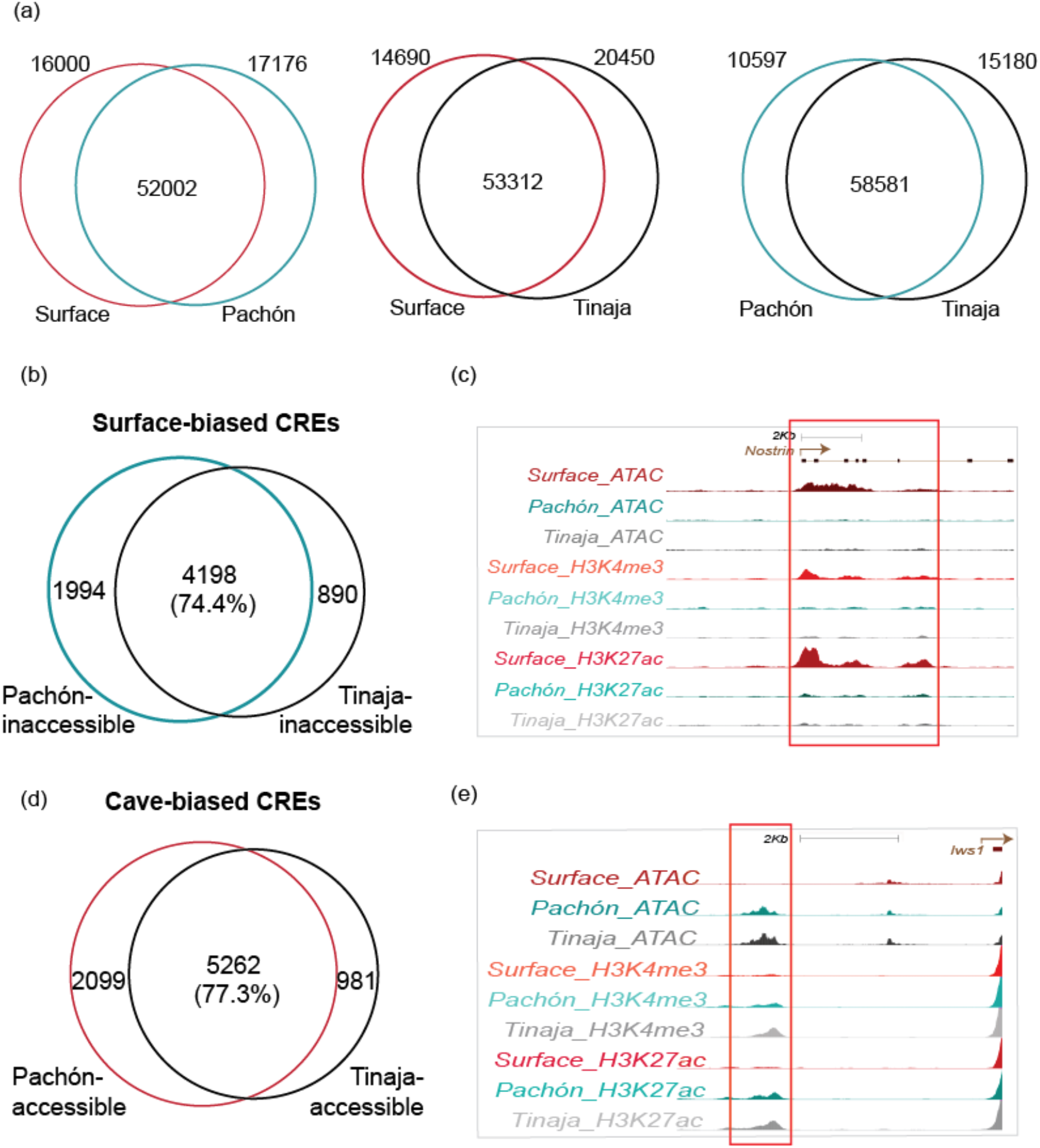
Analysis of morphotype-biased accessible chromatin regions. (a) Venn diagram showing overlap of accessible chromatin regions of surface with Pachón (left), surface with Tinaja (center) and Pachón with Tinaja (right). (b) Venn diagram showing overlap of ATAC peaks between Pachón and Tinaja for surface-biased CREs. (c) Browser shot showing epigenetic landscape for a surface-biased CRE. (d) Venn diagram showing overlap of ATAC peaks between Pachón and Tinaja for cave-biased CREs. (e) Browser shot showing epigenetic landscape for a cave-biased CRE.

Changes in *cis*-regulatory elements often do not show a strict correlation with expression changes of nearby genes because of genomic complexity and redundancy of elements regulating the same loci (Hong et al., 2008; Schmidt et al., 2010; Wong et al., 2015). However, there is evidence that CREs tend to control nearby genes (Fishilevich et al., 2017; Hariprakash and Ferrari, 2019; Mifsud et al., 2015). In addition, studies showed that evolutionarily biased enhancers associated with transcriptional changes are often linked to phenotypic variations (Prescott et al., 2015). Therefore, to focus on stronger CRE-gene correlation, we analyzed CREs within 10kb of the TSS of genes (Daugherty et al., 2017; Hariprakash and Ferrari, 2019; Mifsud et al., 2015). We observed a significant correlation (P<0.05; see methods for details) between morphotype-biased CREs and expression of nearby genes, with s-CREs associated to a larger proportion of surface-upregulated genes and c-CREs associated to more cave-upregulated genes (Fig. 3a. b). We observed the s-CREs associated genes were enriched in circadian clock categories, lipid metabolism, and TGF-β signaling pathways (Fig. 3c), while c-CREs displayed enrichment of multiple pathways involved in lipid metabolism and immune function (Fig.3d). Interestingly, lipid metabolism pathway genes enriched near s-CREs comprised of catabolic genes (lipases and fatty acid binding proteins) while lipid metabolism pathway genes near c-CREs highlighted lipid signaling and anabolic genes (fatty acid synthase and acyl CoA synthetases). These findings are in line with previous studies showing increased fat accumulation in cavefish (Riddle et al., 2018; Xiong et al., 2018).

**Figure 3:**
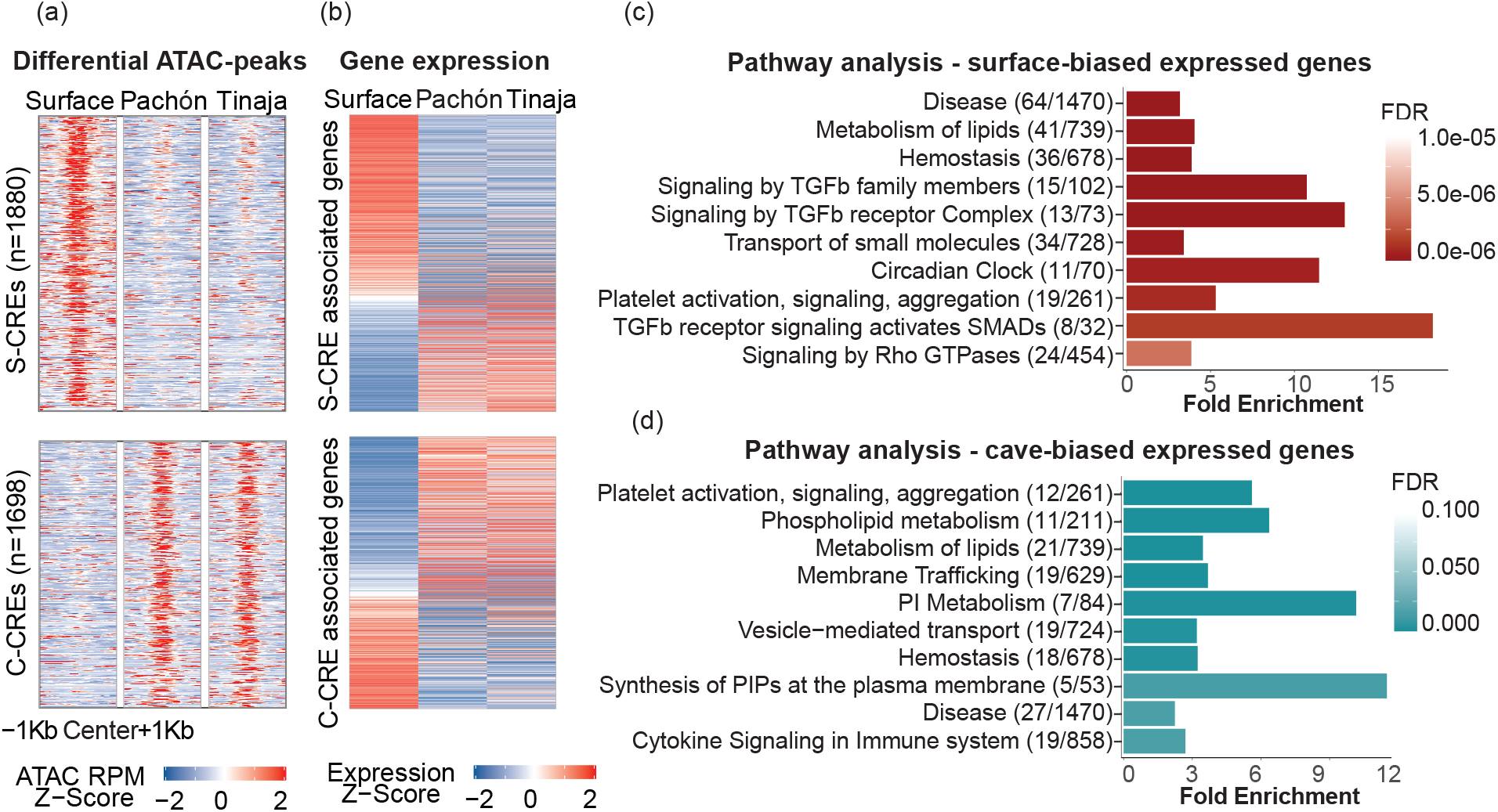
Morphotype-biased accessible chromatin regions associate with key metabolic pathway genes. (a) Heatmaps depicting z-scores of ATAC signal at the s-CREs (top panel) and c-CREs (bottom panel). The number in parentheses indicate the number of peaks in the corresponding panel of the heatmap. (b) Heatmaps depicting z-scores for expression of genes associated to s-CREs and s-CREs. (c, d) Pathways/Reactomes (using GSEA) enriched in genes in proximity (<10kb of TSS) of s-CREs (c) and c-CREs and which show the same bias in expression (d) The figures in parentheses represent the number of genes in our list/ total number of genes in that reactome. The x-axis represents fold change and the color of the bars reflect the FDR value as shown in the color panel on the right side. PI – phosphatidylinositol, PIPs – phosphatidylinositol phosphates.

### Genetic changes are causal for the differential functional output of several CREs

The differential accessibility of the s- and c-CREs could either be due to differences in trans-acting factors or differences in the sequence of the putative CRE itself. To understand the role of specific transcription factors in the differential accessibility, we analyzed enriched binding motifs of transcription factors in s-CREs and c-CREs. In s-CREs, we observed enrichment for motifs of nuclear receptors – Retinoic acid receptor (RXR), Liver X receptor (LXR), and HNF4a, which are known to regulate glucose and lipid metabolism (Fig. S3a) (He et al., 2013; Laurencikiene and Ryden, 2012; Weissglas-Volkov et al., 2006), confirming the results from the pathway analysis (Fig. 3c). Similar analysis for c-CREs revealed enrichment of binding sites for NFY and KLF14 (Fig. S3b). NFY regulates lipid metabolism via the leptin pathway (Lu et al., 2015) and KLF14 represses TGF-β signaling (Truty et al., 2009), a pathway that is enriched in surface fish but not in cavefish (Fig. 3c, d). Notably, we found the consensus motif for CTCF, a factor responsible for regulatory functions including transcriptional activation and repression as well as promoter-enhancer insulation, to be enriched in both s-CREs and c-CREs (Phillips-Cremins et al., 2013). Enrichment of CTCF in these regions supports their putative function as regulatory regions of the genome along with the role of CTCF in the 3D organization of the genome during evolution (Kentepozidou et al., 2020). Together, these analyses point to key liver transcription factors and pathways that likely influence metabolism via the identified CREs.

The binding of transcription factors can directly impact the epigenetic status and function of the CRE (Klemm et al., 2019). We reasoned that the differential accessibility of the s-CREs and c-CREs could be due to differential expression of the transcription factors (TF) that recognize the cognate motifs enriched in the analysis. We searched the liver transcriptome data for these factors (*rar, nfkb, lxr, ctcf, hnf4a, ets, gabpa, c-myc*) and found little or no significant expression difference between morphotypes (Fig. S3c). However, polymorphisms between CREs of surface and cave fish revealed numerous SNPs in TF binding motifs that could negatively affect TF binding (Fig. S3d). This suggests that genetic changes in these putative CREs may contribute to the difference in accessibility.

To functionally validate our predictions, we used a luciferase reporter assay to examine activity of a selected set of surface-biased and cave-biased CREs. To this end, we focused on the differentially accessible CREs that had at least one polymorphism (SNP or indel) between surface and either of the cave populations and had an annotated gene nearby whose expression was biased in the same direction as the chromatin accessibility (Fig. 4a). From these candidates, we evaluated additional parameters such as the degree of difference in the expression of associated gene, maintenance of differential chromatin features in the flanking genomic regions, and relevance of the associated gene in metabolism or liver biology (see methods section for details). Based on these features, we chose a total of 25 differentially accessible CREs for functional validation (Fig. 4a).

**Figure 4:**
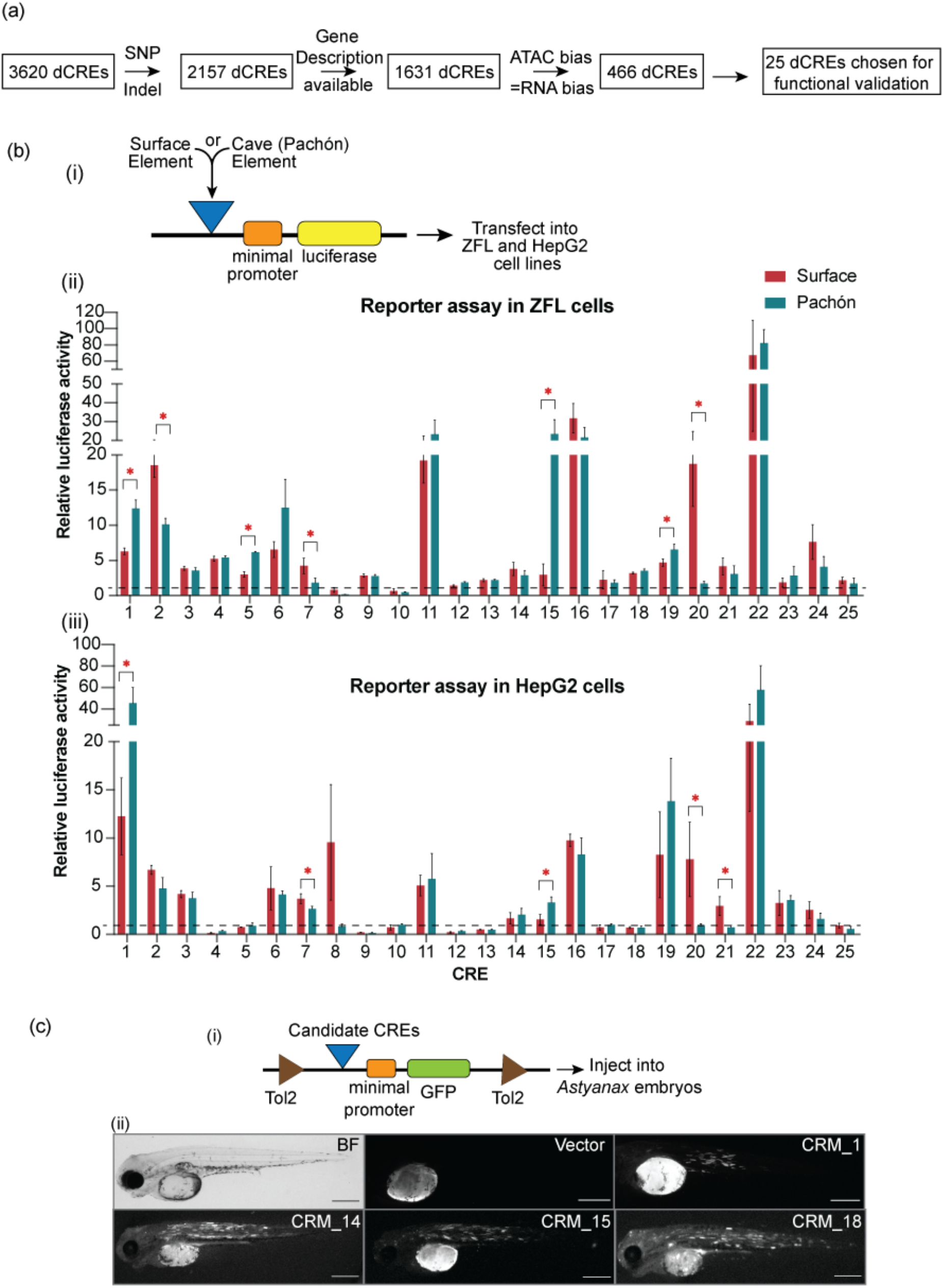
Functional validation of differentially accessible CREs. (a) Flowchart showing the process of selection of candidate differential CREs for functional testing. (b-i) Schematic of the reporter construct used for assaying enhancer activity in cell lines. (b-ii, iii) Enhancer activity for the 25 enhancer candidates each from surface and Pachón tested using luciferase assay in ZFL cells (b-ii) and HepG2 cells (b-iii). * indicates candidates whose enhancer activity is significantly different between surface and Pachón CRE constructs (p-value < 0.05 using two-tailed student’s t-test). Error bars represent standard deviation between 3 biological replicates. The horizontal dashed line marks activity of vector alone (normalized to 1). (c-i) Schematic of the reporter construct used for assaying enhancer activity in *A. mexicanus* larvae. (c-ii) Bright field image of uninjected 4 days post fertilization *A. mexicanus* surface fish larvae (BF), injected with vector alone (Vector) and injected with candidate constructs. Scale bar = 0.5 mm.

In the absence of available *Astyanax* liver cell lines, we chose to perform luciferase reporter assays in the zebrafish liver cell line (ZFL) and the human liver cell line (HepG2). We tested the surface and the Pachón alleles for each of the 25 CREs in replicates. We found that 80% of the tested CREs (20 out of 25) (Fig. 4b-i, ii) mediated expression 2-fold or higher than the empty vector control in ZFL cells, indicating their ability to function as enhancers in cell lines. We observed 32% of these CREs were also functional in human cells (Fig. 4b-iii), suggesting conservation of regulatory function across large evolutionary distances. Additionally, for 7 out of the 20 functional CREs, the surface and cave alleles displayed differential enhancer capabilities when tested in ZFL cells while 5 out of 8 CRE enhancers were differential in HepG2 (Fig. 4b-ii, iii). Combining the results, we observed that CRE_1, CRE_7, CRE_15 and CRE_20 maintained their differential reporter output in both the tested cell lines, reaffirming that polymorphisms in the underlying DNA sequences of these CREs are causal for differences in their ability to drive reporter expression. To further test the CREs’ functionality *in vivo*, we tested 4 out of the 20 enhancer CREs validated in cell lines, in surface fish (Fig. 4c-ii) and zebrafish embryos (Fig. S4). We observed enhanced GFP expression compared to vector control, suggesting robust activity of these enhancers spanning broad developmental timepoints.

### Increased activity of CRE_15 in cavefish is due to deletion of a repressor binding site

We further characterized the cave-biased enhancer CRE_15 (hereafter *E-hpdb*) because of its close proximity (744bp) to the transcription start site of *hpdb* (4-hydroxyphenylpyruvate dioxygenase b) (Fig.5a), a gene that catalyzes the first unidirectional step in tyrosine catabolism. *hpdb* is upregulated in the Tinaja liver with respect to surface (Fig. S5a) and the most upregulated gene in the transcriptome of Pachón (Fig. S5a), which we also validated using qPCR (Fig. 5b).

**Figure 5:**
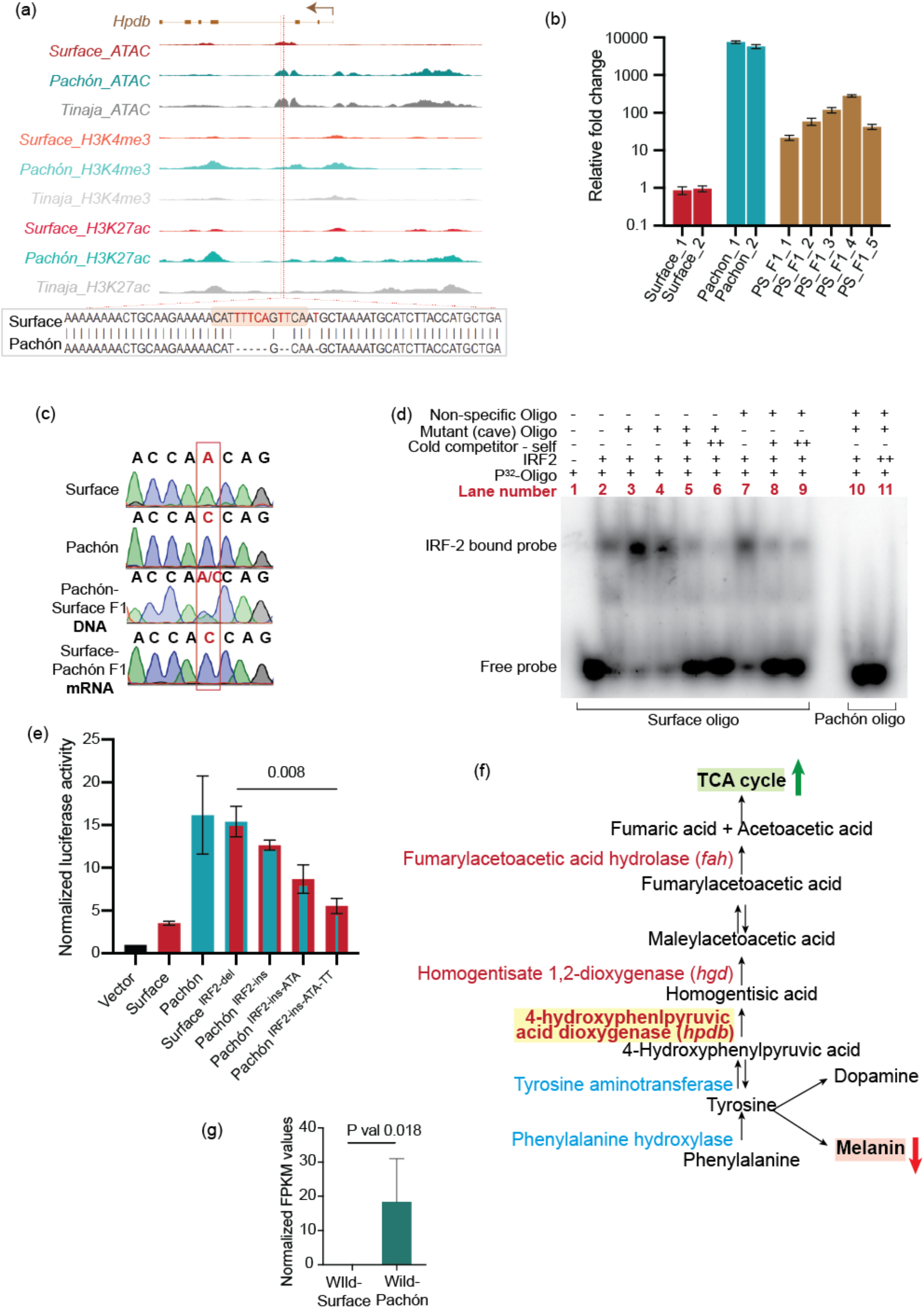
Detailed characterization of CRE_15. (a) UCSC genome browser shot showing various chromatin features at the genomic region around CRE_15. The lower panel shows the sequence at the marked site which includes deletion removing the predicted IRF2 binding site. (b) *hpdb* RNA levels using qPCR in livers of adult surface fish, Pachón cavefish and Pachón-surface F1 hybrids. Each bar represents data from one individual fish. All values are relative to Surface_1 fish normalized to 1. (c) Gel-shift assay for recombinant IRF-2 binding on surface oligo and mutant cave oligo. Excess P-32 labeled oligo runs at the bottom while IRF-2 bound surface oligo runs slower. Cave oligo fails to bind the protein. (d) Relative luciferase activities for vector alone and various alleles of CRE_15 – surface, Pachón, S-CRE_15 without IRF2 binding site (surface^IRF2-del^) and P-CRE_15 with the IRF2 binding site restored (Pachón^IRF2-ins^), Pachón^IRF2-ins^ with additional mutations converting Pachón allele to Surface allele (Pachón^IRF2-ins-ATA^ and Pachón^IRF2-ins-ATA-TT^). (e) Chromatograms from the sequencing of the SNP within exon12 of the *hpdb* gene used to distinguish surface and Pachón alleles. (f) Schematic of pathways which use tyrosine in the cell. A decreased demand for melanin in cavefish could in principle lead to increased availability of tyrosine for other pathways. (g) *hpdb* RNA levels in wild-caught surface fish and Pachón cavefish livers (RNA-seq data from Krishnan et al., 2020).

To genetically assess whether the decreased expression of *hpdb* in surface fish is due to a *cis*-mediated repression or due to the presence of some trans-acting factor(s), we took advantage of the ability to detect allele specific expression in Pachón-surface hybrid F1 crosses (Wittkopp et al., 2004). The qPCR quantification of *hpdb* expression in surface and Pachón matched the RNA-seq data, and the livers of the F1 hybrid fish expressed intermediate levels of the RNA indicating that the Pachón allele is incomplete dominant (Fig. 5b). Next, to test if the expression of *hpdb* RNA is regulated in *cis*, we monitored allele-specific expression levels by taking advantage of the presence of a synonymous SNP in exon 12 of the *hpdb* coding region to distinguish the parental alleles (Fig 5c). While we detected both alleles in the DNA samples, we detected only expression from the Pachón allele in the mRNA samples. This suggests that the increased expression of *hpdb* in Pachón is mediated by changes in *cis*.

We identified multiple small deletions in the *E-hpdb* sequence in Pachón that are predicted to abolish a binding motif for the repressor protein IRF2 (Fig. 5a, lower panel). As there was no significant difference between *irf2* expression between surface and Pachón (Fig. S5b), we explored the possibility that differential binding of IRF2 protein could be linked with the regulatory difference in *E-hpdb* by using an *in-vitro* electromobility shift assay (EMSA) (Fig. 5d). We carried out EMSA using γP-32 labeled 20bp oligonucleotides from the surface fish enhancer containing the IRF2 binding site and the corresponding 20bp region from the Pachón enhancer lacking that site. We used recombinant human IRF2 protein as the DNA-binding domain is highly conserved with that of the *Astyanax* Irf2 protein and identical between Pachón and surface fish (Fig. S5c). IRF2 binds to the oligo based on the surface fish sequence, while very weak or no binding was observed with the Pachón probe, showing altered affinity of this region for the IRF2 protein (Fig 5d). We also tested the specificity of IRF2 binding on the surface *E-hpdb* by using Pachón *E-hpdb* and unrelated oligos as non-specific competitors and observed a robust DNA-protein interaction (Fig. 5d). In addition, we performed cold competition by the addition of 200X (lane-5 and 8) and 400X (lanes-6 and 9) unlabeled self-competitor and observed weakening of the interaction, confirming the specificity of the binding. These results suggest that the deletion in P*-E-hpdb* prevents the IRF2 repressor from binding, which in turn could have an effect on the activity of this regulatory region.

We next tested if the lack of an IRF2 site is sufficient to abolish repression *in vitro*. Using the cell-based luciferase assay, we found that specifically deleting the IRF2 motif in the surface allele of *E-hpdb* (Surface^IRF2-del^ – now similar to the Pachón allele) restored expression (Fig. 5e). This suggests that IRF2 is indeed a repressor of *E-hpdb*. Next, we asked whether repression can be rescued by adding back the IRF2 binding site to the Pachón sequence. Surprisingly, we noted that the addition of the IRF2 site was not sufficient to cause significant repression (Pachón^IRF2-ins^) (Fig. 5e), suggesting that other sequence changes have occurred in Pachón that prevent repression by IRF2. We searched for other mutations and sequentially edited two more sites (Pachón^IRF2-ins-ATA^ and Pachón^IRF2-ins-ATA-TT^) by converting them from the surface alleles to the Pachón alleles. When tested in the luciferase assay, we observed that the additional mutations restored the repressive activity of the IRF2 site insertion in P-*E-hpdb*. This suggests that while deleting the IRF2 binding site is sufficient to release repression, other mutations have contributed to the differential activity of the CRE.

### Deletion of IRF2 binding site in independent cave populations as an adaptive trait

Tyrosine serves as a substrate for several important compounds in the cell, including melanin, dopamine and certain intermediates for ketone body formation and the TCA cycle (Fig. 5f). An accumulation of excess tyrosine in cells has been reported in cave populations that are mutant for melanin formation and in surface fish upon abrogation of melanin formation by knocking down *oca2* (Bilandzija et al., 2013). We also noticed in our liver transcriptome data that the expression of genes encoding enzymes that convert tyrosine to L-Dopa (Tyrosinase and tyrosine hydroxylase) is very low (Fig. S5d). In a nutrient deprived condition, it could be a thrifty strategy to divert the excess tyrosine to produce TCA intermediates and ketone bodies for energy storage and production.

We investigated this possibility by analyzing the deletion of the IRF2 binding site and the difference in the *hpdb* expression in wild surface and independent cave populations. We genotyped 23 wild-caught surface fish and 23 wild Pachón fish and found that the IRF2 site deletion was fixed in all of the sequenced Pachón fishes, while all surface fishes had the wildtype sequence. This was in line with our published data wherein the expression of *hpdb* from wild Pachón was higher than wild surface fish (Fig. 5g) (Krishnan et al., 2020). To extend this analysis we also genotyped other cavefish populations (Tinaja, Yerbaniz, Piedras, and Japonés) and found the same deletion to be fixed in all populations, except for the Japonés individual, which was heterozygous. These observations suggest that the mutation is under positive selection (Bilandzija et al., 2013; Chan et al., 2010). We further analyzed the expression of other relevant enzymes in the tyrosine catabolism pathway and found that genes encoding enzymes *hpdb*, *hgd* and *fah*, that catalyze unidirectional steps in the tyrosine catabolism pathway, are expressed higher in Pachón than in surface livers (Fig. 5b, Fig. S7e). This supports our hypothesis that excess tyrosine in the cells is being repurposed via the TCA cycle.

## Conclusions

In this study, we utilized *Astyanax mexicanus*, an evolutionary and genetic model system to address the question of how changes in the cis-regulatory elements of the genome, can help the cavefish adapt to the extreme cave environment characterized by absence of light and nutrient deprivation. Our analysis uncovered a large number of putative cis-regulatory elements that are differentially regulated between the surface and the cave morphotypes. We found surface-biased CREs to be associated with key genes related to lipid catabolism, circadian rhythm and the TGF-β signaling, while cave-biased CREs were associated mainly with lipid anabolism genes and immune system. The biased enrichment of motifs for key transcription factors like Liver X Receptor and HNF4a in s-CREs, and NFY and KLF14 in c-CREs points to key transcription factor networks that could be linked to changes in the above metabolic pathways. These global regulatory changes reveal how adaptation to the cave environment has modified the regulatory architecture of the genome to support physiological traits and metabolic processes in cavefish compared to surface fish

Our study also highlights that the genome-wide chromatin architecture of the two cavefish morphotypes – Pachón and Tinaja – were more similar to each other than to the surface fish. This is in line with the previously observed phenotypic convergence in these independently derived cavefish populations like loss of visible eyes and pigment, accumulation of excess fat, insulin resistance, etc. (Jeffery, 2009; Riddle et al., 2018; Xiong et al., 2018). To what extent this stems from standing genetic variation, phylogenetic relationship, or common upstream modulators remains to be studied further, however, one interesting hypothesis is that selection has repeatedly acted on existing transcriptional gene regulatory networks in the animals upon exposure to new environments.

Functional validation of our genome-wide analyses in cell culture assays revealed that a large proportion of CREs have altered activities potentially due to trans-acting effects while some arise due to underlying genetic changes in them. Detailed genetic and functional characterization of an enhancer of the *hpdb* gene affirmed the role of cis-regulatory changes in controlling the cave-biased expression of *hpdb* gene in the liver. The *hpdb* gene is part of the tyrosine metabolism pathway and converts tyrosine to TCA cycle intermediate fumarate and ketone body acetoacetate. It has been shown that in the absence of melanin, as seen in cavefish, results in more tyrosine and dopamine in the body (Bilandzija et al., 2013). An intriguing possibility supported by the presence of this mutated CRE in multiple cave populations is that, in the liver, excess tyrosine can be converted to fumarate and acetoacetate as a measure to use any available nutrient for energy production by cavefish.

We expect many other differentially accessible regions identified in our study to be involved in the metabolic adaptation of cavefish to their low-nutrient environment. We propose that *Astyanax mexicanus,* with its contrasting morphotypes and independently derived cave populations, presents an effective system to unravel global gene regulatory pathways and networks important in physiological adaptation of species to new and changing environments that could give insights into a better understanding of conserved metabolic processes in vertebrate physiology.

## Methods

### Astyanax husbandry

*Astyanax* are housed in polycarbonate or glass fish tanks on racks (Pentair, Apopka, FL) with a 14:10 h light:dark photoperiod. Each rack uses an independent recirculating aquaculture system with mechanical, chemical, and biologic filtration, and UV disinfection. Water quality parameters are maintained within safe limits (upper limit of total ammonia nitrogen range 1 mg/L; upper limit of nitrite range 0.5 mg/L; upper limit of nitrate range 60 mg/L; temperature set-point of 22°C; pH 7.65, specific conductance 800 μS/cm; dissolved oxygen >90%. Water changes range from 20-30 % daily (supplemented with Instant Ocean Sea Salt [Blacksburg, VA]). Adult fish are fed three times a day during breeding weeks and once per day during non-breeding weeks on a diet of Mysis shrimp (Hikari Sales USA, Inc., Hayward, CA) and Gemma 800 (Skretting USA, Tooele, UT).

### Zebrafish husbandry

Zebrafish are housed in polycarbonate fish tanks on racks (Pentair Aquatic Eco-Systems, Inc., Apopka, FL) with a 14:10 h light:dark photoperiod. Racks are supplied by two recirculating aquaculture systems with mechanical, chemical, biological filtration, and UV disinfection. Water quality parameters are maintained within safe limits (upper limit of total ammonia nitrogen range 0.5 mg/L; upper limit of nitrite range 0.5 mg/L; upper limit of nitrate range 40 mg/L; temperature set-point 28.5°C; pH 7.60, specific conductance 500 μS/cm; dissolved oxygen >85%. Water changes range from 20-30 % daily (supplemented with Instant Ocean Sea Salt [Spectrum Brands, Inc., Blacksburg, VA]). Adult zebrafish are fed twice daily with one feed of hatched Artemia (1^st^ instar) (Brine Shrimp Direct, Inc., Ogden, UT) and one feed of Zeigler Adult Diet (Zeigler Bros, Inc., Gardners, PA. Embryos up to 5dpf were maintained at 28.5°C in E2 embryo media.

### ATAC-seq

Livers were dissected from 3 fish each for surface, Pachón and Tinaja populations and divided into two parts for RNA-seq and ATAC-seq. ATAC-seq was performed as per Buenrostro et al. (Buenrostro et al., 2015) with some modifications to accommodate the use of whole tissues as starting material instead of cells. Livers were homogenized (30-40 strokes) using the loose pestle of Dounce homogenizer in lysis buffer (10 mM Tris·Cl, pH 7.4, 10 mM NaCl, 3 mMMgCl2, 0.1% (v/v) Igepal CA-630) to prepare nuclei. Rupturing of cell membrane and obtainment of nuclei was confirmed using Trypan blue staining under a phase contrast microscope. Nuclei were counted under microscope and ~70000 nuclei were taken and spun down at 1500g at 4 °C. The nuclei were resuspended in transposition mixture (25 μl TD of 2× reaction buffer from Nextera kit, 2.5 μl TDE1 Tn5 Transposase from Nextera kit, 22.5 μl nuclease-free water) and incubated at 37 °C for 22-24min. The reaction was purified, and the library was prepared as per the original protocol (Buenrostro et al., 2015) Paired-end sequencing was performed using the Illumina Next-seq Mid-output mode.

### RNA-seq

The part of the livers that was used for RNA-seq was frozen and later used for RNA extraction using the Qiagen RNeasy kit. RNA-seq and ATAC-seq were performed using the same liver samples in order to get maximum possible correlation between chromatin accessibility. Libraries were prepared according to manufacturer’s instructions using the TruSeq Stranded mRNA Prep Kit (Illumina). The resulting libraries were purified using the Agencourt AMPure XP system (Beckman Coulter) then quantified using a Bioanalyzer (Agilent Technologies) and a Qubit fluorometer (Life Technologies). Libraries were re-quantified, normalized, pooled and sequenced on an Illumina HiSeq 2500 instrument as 50-bp single read. Following sequencing, Illumina Real Time Analysis version 1.18.64 and bcl2fastq2 v2.20 were run to demultiplex reads and generate FASTQ files.

### Chromatin Immunoprecipitation

Adult livers were dissected and fixed in 1% formaldehyde for 15 min. Chromatin was prepared using the MAGnify Low Cell ChIP-Seq Kit (Thermo Fischer) and ChIP was done for H3K4me3 (Millipore, # 07-473), H3K27ac (Abcam, # ab4729), H3K27me3 (Abcam, # ab195477). 200,000 cells and 1ug of antibody was used for each sample. Libraries were prepared using the KAPA HTP Library Prep Kit for Illumina and Bioo Scientific NEXTflex DNA barcodes. The resulting libraries were purified using the Agencourt AMPure XP system (Beckman Coulter) then quantified using a Bioanalyzer (Agilent Technologies) and a Qubit fluorometer (Life Technologies). Post amplification size selection was performed on all libraries using a Pippin Prep (Sage Science). Libraries were re-quantified, normalized, pooled and sequenced on Illumina HiSeq 2500 instrument as 50-bp single read. Following sequencing, Illumina Real Time Analysis version 1.18.64 and bcl2fastq2 v2.18.0.12 were run to demultiplex reads and generate FASTQ files.

### Analysis

FASTQ files for each chromatin mark were aligned to astMex_2.0 using bowtie2 under default parameters. Reads mapping to the mitochondrial chromosome were subsequently filtered out using samtools prior to calling peaks with MACS2 under default parameters. ATAC-seq data was aligned to astMex_2.0 with bowtie2 with the following parameters: −X 2000 ‒very-sensitive. Peaks were called with MACS2 using default parameters. For each chromatin mark and ATAC peak, a reference set of peaks was created for each type of fish by reducing peaks from each replicate to a single peak set, and then keeping only peaks observed in at least two replicates. These loci were quantified and analyzed for differential openness between fish using edgeR to calculate logFC ratios and p-values. ATAC-seq loci were then scored for overlap (1 or 0) with each chromatin mark and mapped to the nearest gene using the GenomicRanges library in R. Peaks were filtered for logCPM values greater than 0. For differential peaks, the p-value for the difference in the peak intensities between 2 morphotypes was <0.0001. For analysis of ChromHMM, pathway and motif enrichment we used CREs within 10kb of the nearest TSS.

### Heatmaps

Heatmaps for signals around transcription start sites (TSS) i.e. promoters show the reads per million (RPM) normalized to ChIP-seq signals around TSS regions (2 Kb upstream and downstream) of 18836 transcripts. Transcripts were selected by 1. Ensembl 94 protein-coding genes; 2. Longest transcript for each gene; 3. No overlapping transcripts. The 4 Kb TSS regions were binned into 100 bins (40 bp per bin) and average RPM signal was calculated for each bin. TSS regions were ordered was based on average ATAC signals in decreasing order.

Heatmaps for differentially accessible CREs consist of 1698 cave-biased and 1880 surface-biased peaks with q-value < 0.05. In addition, the differential peaks had q-values <0.05 for both Pachón vs surface and Tinaja vs surface. Conversely, invariant peaks represented in the heatmap had q-values (Pachón vs. surface, Tinaja vs. surface, and Pachón vs. Tinaja) were all greater than 0.90 (1991 peaks). We plotted the reads per million (RPM) normalized ATAC signals (row scaled as Z-scores) in ±1Kb regions around peak centers using R package EnrichedHeatmap (Gu et al., 2018). We plotted the expression (normalized TPM, row-scaled) of the nearest gene (within 10Kb region) for differentially accessible CREs in the three fish types. Genes are ordered based on average fold change between surface fish and cavefish. For surface-biased peaks, genes express higher in surface fish than cavefish (one-sided t-test p-values are 0.014 for surface vs. Pachón and 0.052 for surface vs. Tinaja); and for cave-biased peaks, genes express higher in cavefish than surface fish (one-sided t-test p-values are 0.033 for Pachón vs. surface and 0.031 for Tinaja vs. surface).

### Conservation

All conservation analyses were done using the AstMex1 genome. Genomic Evolutionary Rate Profiling (GERP) regions and their conservation scores for 11 fish conservation were obtained from Ensembl. To obtain evolutionarily conserved CREs, overlap was seen between open chromatin regions and evolutionarily constrained regions from GERP. We omitted exons from the GERP regions in order to prevent bias from the highly conserved exonic regions. To obtain the background level of conservation, size-matched random set of regions were obtained from the genome and the process was iterated 1000 times.

To find orthologues of known human liver enhancers, human multiz100way genome alignments were downloaded from UCSC and pairwise alignments between human (hg19) and cavefish (astMex1) were extracted using UCSC utilities (mafSpeciesSubset). Then bedtools (v2.26.0) was used to map human enhancers to their nearest conserved regions (Quinlan and Hall, 2010).

### ChromHMM

We applied ChromHMM (v1.15) on H3K27ac, H3K4me3, and H3K27me3 histone marks for three fish types (surface, Pachón, and Tinaja). Cavefish genome for ChromHMM was built using Ensembl 94 cavefish genome and annotation release. ChIP-seq BAM files were binarized using BinarizeBam command with bin size (−b option) of 200 and Poisson threshold (−p option) of 0.001. Next we built a 7-state hidden Markov model (HMM) using LearnModel with default parameters. ChromHMM State Distribution: ATAC consensus peaks were resized to 400 bp around peak center. Cavefish genome was binned into 200 bp non-overlapping bins using bedtools makewindows command. We then assigned a ChromHMM state to each ATAC peak or genome bin, requiring the state covered at least 50% of the ATAC peak or genome bin. Next ChromHMM state distribution was calculated for ATAC peaks and genome bins by counting the occurrences of each assigned state.

ChromHMM State Enrichment: We randomly placed the ATAC consensus peaks (resized to 400 bp) onto cavefish genome using bedtools shuffle (with -noOverlapping option) command. This shuffling process was repeated for 1000 times. For each shuffled ATAC peaks, we computed its ChromHMM state distribution in the same way mentioned above and calculated the log2 enrichment ratio between the true and shuffled distribution. The barplot showed the average enrichment ratio and the standard deviation (n=1000) for each ChromHMM state.

### Motif and pathway analysis

Motif enrichment analysis was done using HOMER (Heinz et al., 2010) with the command ‘findMotifsGenome.pl - size given’ and known vertebrate motifs were analyzed. Pathway (Reactome) analysis was performed using Molecular Signatures Database of the Gene Set Enrichment Analysis (Mootha et al., 2003; Subramanian et al., 2005) with default parameters.

### motifBreakR

Single nucleotide polymorphisms in cavefish and surface fish were annotated for their potential effect on transcription factor binding motifs using the R package motifBreakR [1]. The main function of the package was run with the following parameters: snpList (an R object to read directly from the VCF file of cave and surface SNVs), filterp=TRUE, pwmList = homer_motifs (motifs from HOMER database), threshold=1e-4, method=”ic”, and bkg=c(A=0.25, C=0.25, G=0.25, T=0.25). The resulting GRanges object containing SNP effect on transcription factor binding motifs was used for downstream analysis.

### SNP calling for selecting polymorphic CREs

Reads from FASTQ files were aligned with BWA mem and marked by Read Group. BAM files were merged and deduplicated using GATK. Reads were processed for calling variants with HaplotypeCaller using a GATK best practices pipeline implemented with Snakemake. Two rounds of boostrapping with filtering were used to create a reference set of variants for Base Quality Score Recalibration prior to calling variants against astMex_2.0

### Selection of candidates for functional testing

We set a series of criteria by which we selected differential CRE candidates for functional testing. After selecting for candidates with polymorphisms, a well annotated neighboring gene and a biased gene expression, we were left with 466 candidates to choose from. We first looked at the details of the genomic context of the CRE. We checked if the flanking regions of the CRE maintained the biased epigenetic signature and that there were no major unbiased peaks in the immediate flanking regions. We next looked for various characteristics of the neighboring gene. We selected CREs that were associated to genes with highly differential expression levels in our RNA-seq analysis for example, *Nos* and *Hpdb*. Lastly, we reviewed literature and focused on CREs associated with genes that were involved in metabolic processes like carbohydrate or fat metabolism or pathways that maintain health of the tissues like inflammatory pathway, etc. Supplementary Information SI-3 lists the final list of candidates tested in reporter assays along with details of the epigenetic signature and the associated genes.

### Cloning and reporter assays

Candidate CREs from surface, Pachón and Tinaja genomes were amplified from genome or synthesized commercially (GenScript) and cloned into pGL4.23 (Promega) and HLC (Parker et al., 2014) vectors using Gibson assembly (NEB, Cat # E2611). All primers used in the study along with their descriptions are listed in Supplementary Information SI-4.

Reporter assays were performed in *Astyanax* surface fish embryos and zebrafish embryos, adult zebrafish liver cell line ZFL and human liver cell line HepG2. For zebrafish microinjections, differential CRE candidates were cloned into a Tol2-based vector with minimal c-*fos* promoter and downstream GFP. *Astyanax* and zebrafish larvae were anesthetized using buffered MS-222 and immersed in 3% methyl cellulose in a depression slide and imaged using Leica Stereomicroscope.

For luciferase reporter assays, the CRE candidates were cloned into the pGL4.23 vector (Promega) at the EcoRV site upstream to a minimal promoter driving firefly luciferase gene. To control for transfection efficiency, a pRLTK vector (Promega) that expressed Renilla luciferase gene was co-transfected. All transfections were done using Lipofectamine LTX with Plus reagent (Cat # 15338030) in 24-well plates with 350ng of test construct and 150ng of control plasmid. Luciferase activity was measured 48 hours post transfection using a luminometer (Victor X Light, Perkin Elmer). All constructs were done in 2 or more replicates and relative enhancer activity was calculated by normalizing empty vector to 1. Significance was determined using two-tailed Student’s t-test.

### Electromobility shift assays

Gel-shift assays were performed using recombinant Human IRF2 protein (Sigma, #SRP6338). 10 pmoles of single-stranded oligos for surface fish IRF2 binding site and corresponding mutant cavefish IRF2 site were end-labeled with P32 radioisotope using polynucleotide kinase, annealed with complementary strands to make double-stranded oligos and purified using G-25 spin columns (GE Healthcare, #27-5325-01). Binding reactions were set up as follows: 1x binding buffer (5x Binding buffer: 50mM Tris 7.5, 375mM NaCl, 5mM EDTA, 30% Glycerol, 15mM Spermidine, 5mM DTT), 50 ng/μL polydI•dC, 0.25ug protein, 100 fmol of labeled target probe and 25pmol of mutant or non-specific oligo in a final volume of 20 μL. The binding reaction was set up in cold and then incubated at room temperature for 20 min. DNA-protein complexes were separated on non-denaturing 6% DNA retardation gels (Invitrogen, #EC63655BOX) at 100V constant voltage for 50 min. Post-run, the gel was fixed in gel fixation solution (40% v/v Methanol, 10% v/v Acetic acid) for 30min and exposed to Phosphor Imaging screen overnight and imaged using Typhoon scanner.

## Acknowledgments

We are grateful to the cavefish and aquatics core facilities at the Stowers Institute for support and husbandry of the cavefish and zebrafish. DNA samples for wild-caught Tinaja, Yerbaniz, Piedras and Japonés cavefish were generously provided by Richard Borowsky, and Bill Jeffery provided *Astyanax* liver samples for preliminary ChIP experiments. We thank Malcolm Cook for help with motif analysis software, Kyle Weaver for helping with high-throughput genotyping and Mark Miller for illustrations. We thank Robb Krumlauf, Narendra Singh and Julia Zeitlinger for useful inputs throughout the study and critical reading of the manuscript. NR is supported by institutional funding, funding from the JDRF, the Edward Mallinckrodt Foundation, NIH Grant R01 GM127872 and NSF IOS-1933428 and EDGE award 1923372. R.P. was supported by a grant (no. PE 2807/1-1) from Deutsche Forschungsgemeinschaft.

## Author contributions

JK and NRO designed the study. JK performed experiments with support from RP, AK, SX and JWC. RP and AK collected wild *Astyanax* samples. CWS, NZ, JK, JVC performed the analyses with support from HL. JK and NRO wrote the manuscript. All authors read and approved of the manuscript.

## Data availability

Original data underlying this manuscript can be accessed from the Stowers Original Data Repository at http://www.stowers.org/research/publications/libpb-1538. The ATAC-seq, ChIP-seq and RNA-seq data can be found at GEO accession number GSE153052.

