## Supplementary Figures for "Genome-wide analysis of *cis*-regulatory changes in the metabolic adaptation of cavefish"

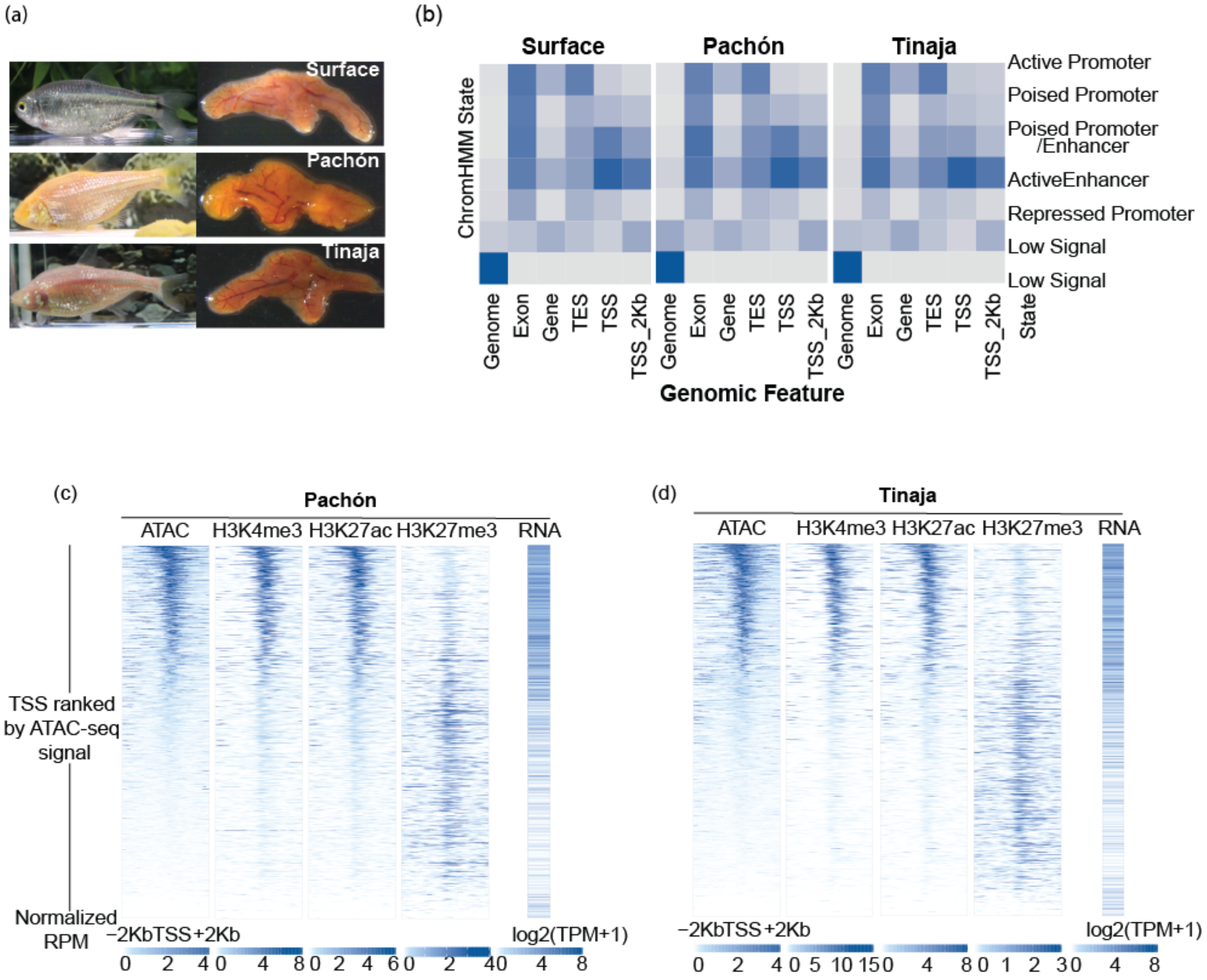


Figure S1: (a) Left panel shows representative images of surface, Pachón and Tinaja fishes and right panel shows their corresponding livers. (b) ChromHMM emission heatmap showing the distribution of various chromatin states across various features in the genome of Pachón and Tinaja. TES – Transcription end sites, TSS – Transcription start sites. (c, d) Heatmaps for ATAC-seq, H3K4me3, H3K27ac and H3K27me3 signal at all the promoters (TSS ±2Kb) along with the corresponding RNA levels for the Pachón and Tinaja morphotypes. Plotted are reads per million (RPM) normalized to ChIP-seq signals at promoters

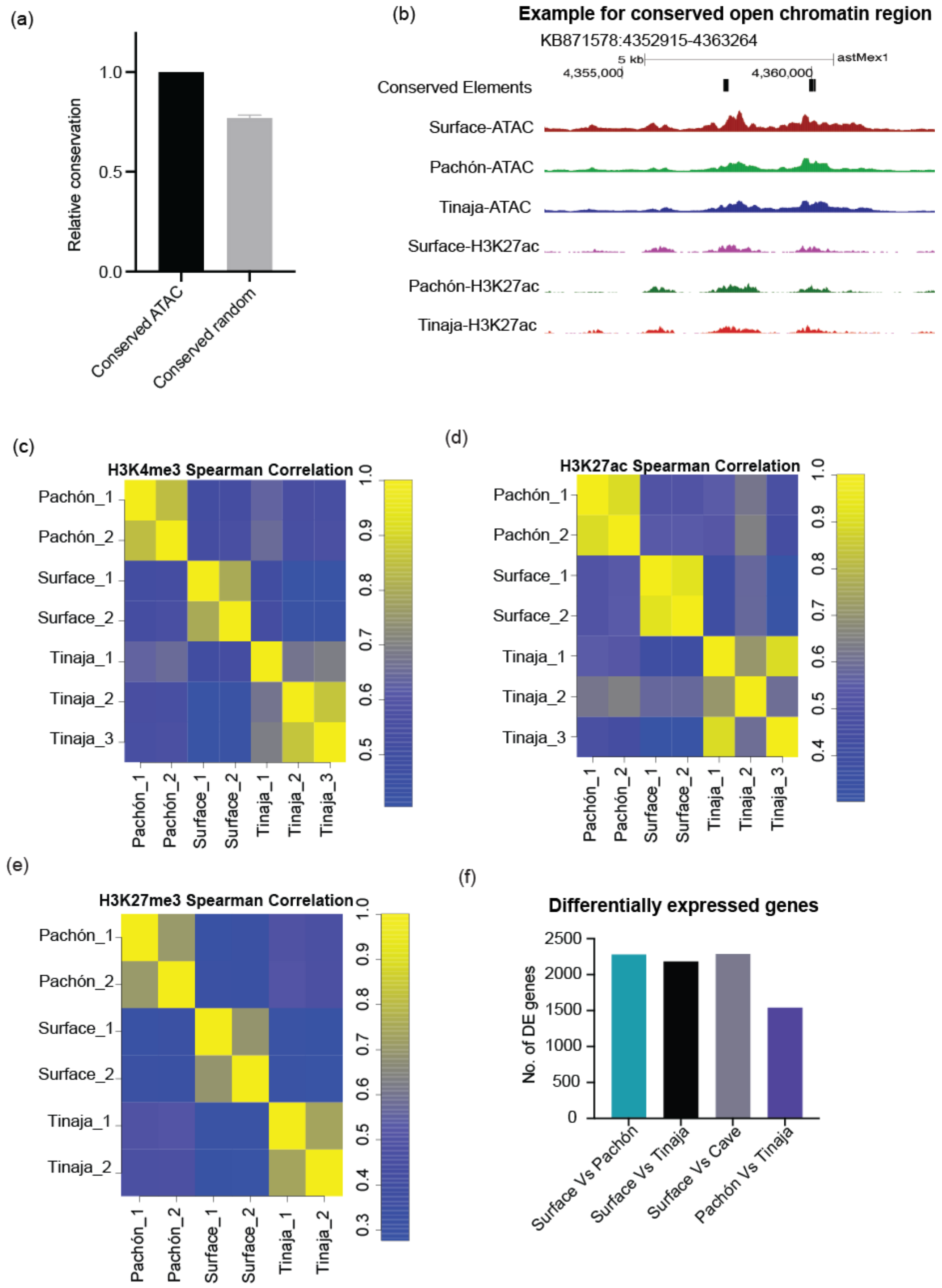


Figure S2: (a) Overlap of ATAC peaks with evolutionarily conserved regions using 11-fish GERP scores normalized to 1 (Black bar). The grey bar represents relative conservation of same number of randomly picked, non-exonic and size-matched genomic regions. The error bar represents standard deviation from 1000 iterations of picking random set of regions (b) Browser shot of one of the conserved candidate CREs. (c-e) Heatmaps for spearman correlations between different samples for each chromatin feature measured in the study – H3K4me3 (c), H3K27ac (d), H3K27me3 (e). (f) Number of differentially expressed genes between different morphotypes.

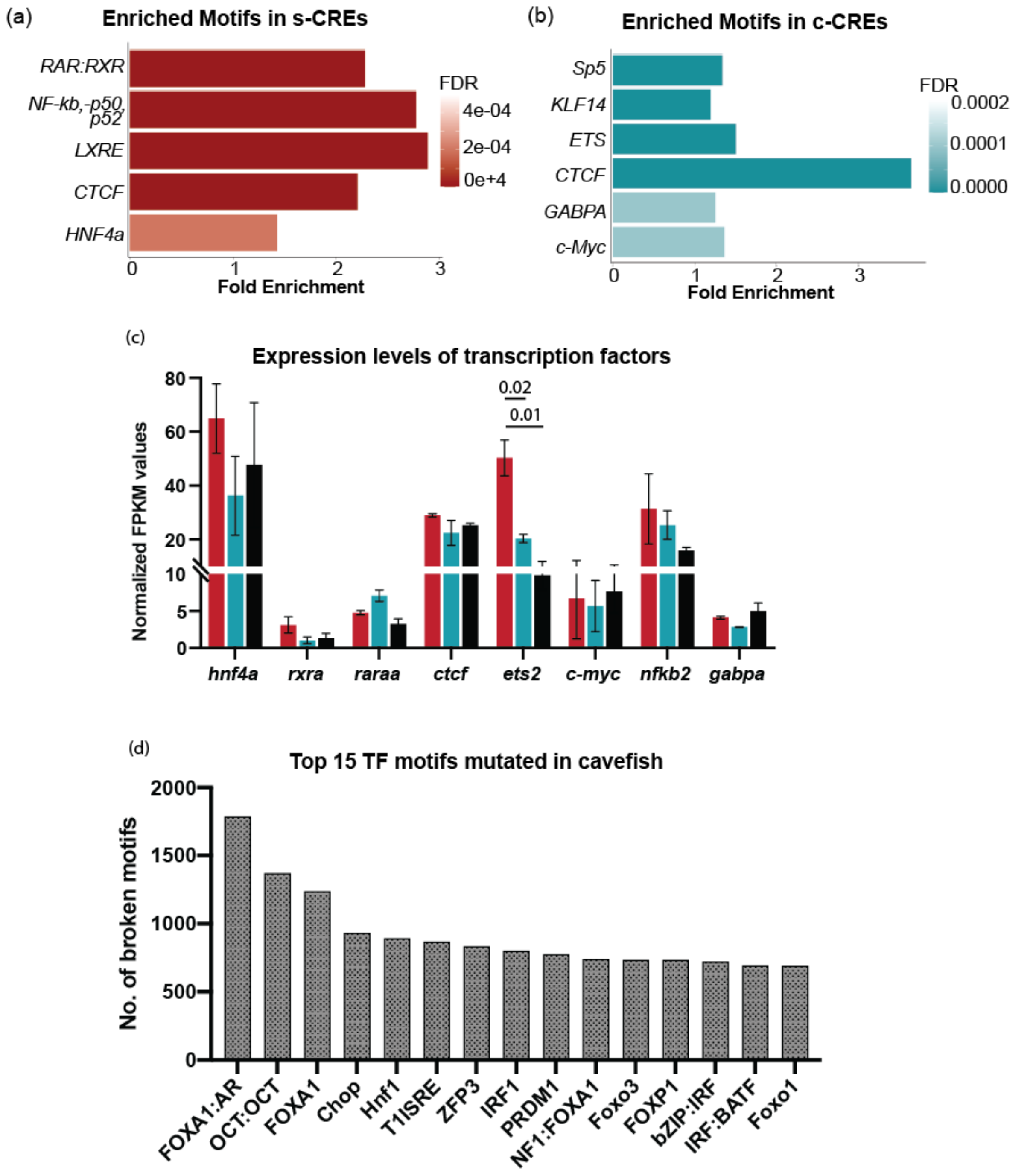


Figure S3: (a, b) Motifs enriched in s-CREs (a) and c-CREs (b). The X-axis represents fold change and the color of the bars reflect the FDR value as shown in the color panel on the right side. (c) Expression levels of transcription factors represented as normalized FPKM values in each morphotype. (d) Top 15 TFs having high numbers of motifs mutated.

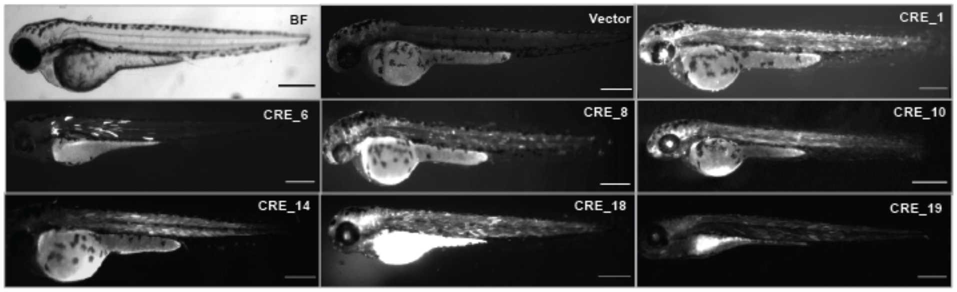


Figure S4: Zebrafish larvae injected with various CRE constructs. On the left top is bright field image of a representative 4dpf zebrafish larva (BF), ‘Vector’ indicates larva injected with vector alone and rest are larvae injected with various candidate constructs. Scale bar = 0.5mm.

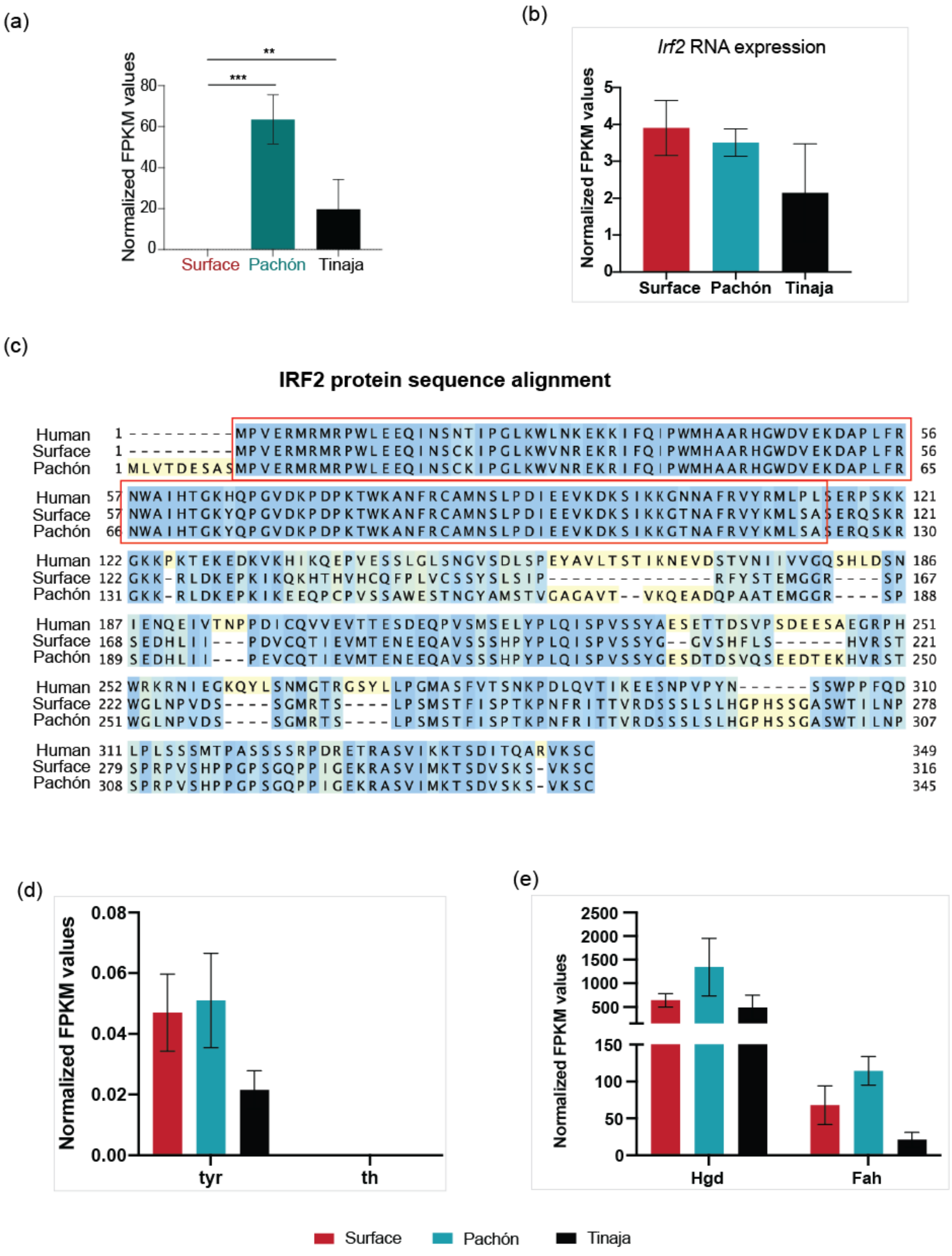


Figure S5: (a) Expression levels of *Hpdb* gene in surface, Pachón and Tinaja livers from RNA-seq experiment. (b) Expression levels of *Irf2* gene in the livers of adult surface, Pachón and Tinaja. (c) Alignment for human and Pachón cavefish IRF2 protein. Amino acids within the yellow box comprises the highly conserved DNA-binding domain of IRF2. (d) Expression levels of genes belonging to tyrosine metabolism that make melanin. Tyr: Tyrosinase, th: Tyrosine hydroxylase. (e) Expression levels of genes belonging to tyrosine metabolism downstream to *Hpdb,* measured in the livers of adult surface, Pachón and Tinaja Hgd: Homogentisate 1,2-dioxygenase, Fah: fumarylacetoacetate hydrolase.
